# Genome-wide survey of tandem repeats by nanopore sequencing shows that disease-associated repeats are more polymorphic in the general population

**DOI:** 10.1101/2019.12.19.883389

**Authors:** Satomi Mitsuhashi, Martin C Frith, Naomichi Matsumoto

## Abstract

Tandem repeats are highly mutable and contribute to the development of human disease by a variety of mechanisms. However, it is difficult to predict which tandem repeats may cause a disease. We performed a genome-wide survey of the millions of human tandem repeats using long read genome sequencing data from 16 humans. We found that known Mendelian disease-causing or disease-associated repeats, especially coding CAG and 5’UTR GGC repeats, are relatively long and polymorphic in the general population. This method, especially if used in GWAS, may indicate possible new candidates of pathogenic or biologically important tandem repeats in human genomes.

## Main text

There are more than 30 rare Mendelian human diseases caused by tandem repeat expansions in human genomes [1]. Genome-wide surveys of tandem repeats in individual genomes are now feasible due to the development of high-throughput sequencing technologies, which enable direct identification of large pathogenic expansions. However, it is still difficult to predict which tandem repeats cause disease, because there are thousands of tandem repeats in each individual that are different from the reference genome. We hypothesize that disease-causing tandem repeats are unstable in the general population. It is unclear whether highly polymorphic repeats cause disease, or stable repeats suddenly expand dynamically in gametes to pathogenic lengths (usually pathogenic expansions are +100 to ∼10,000 repeat units) [2]. Here, we characterize disease-causing tandem repeats using long DNA reads, by measuring the variation of their length, and comparing them to other repeats. Current long read sequencing technologies have achieved reads longer than 10 kb on average, which have a high chance to cover whole tandem repeats including flanking unique sequences. Our recently developed tool, `tandem-genotypes`, can robustly detect tandem repeat changes from whole genome long read sequencing data [3]. We used this tool to genotype tandem repeats based on long reads from nanopore sequencers.

First, we identified tandem repeats in a human reference genome (hg38) using `tantan` [4] (http://cbrc3.cbrc.jp/~martin/tantan/). In total, 3,312,291 loci were identified, with the repeat units ranging from 1 to 100 bp. We used long read whole genome sequencing from 16 humans who do not have any disease or have diseases explained by chromosomal rearrangements (we suppose they do not have pathogenic tandem repeats), with average coverage of x25 (ranging x12-x39, Table S1). `tandem-genotypes` predicted lengths for more than 98% of the 3 million tandem repeats (Table S1, Figure S1), including 215,561 triplet repeats.

Until recently, most of the known disease-causing tandem repeats are CAG or GGC triplet repeats, although there are a few exceptions; quadruplet repeat (GGCT) in Myotonic Dystrophy type 2 (MIM#602668), and sextuplet repeat (GGGGCC) in Frontotemporal dementia and/or amyotrophic lateral sclerosis (MIM#614260)). CAG and GGC triplet diseases have three major disease mechanisms: poly-glutamine diseases (CAG), poly-alanine diseases (GGC), or 5’UTR GGC expansion diseases [5-7]. We investigated 20 CAG and GGC triplet repeat disease loci (Table S2). Distributions of tandem repeat length in all reads from 16 individuals show that disease-causing repeats have more variation than other repeats (the same number of repeat loci were randomly extracted from non-disease repeat loci for comparison to the disease repeat loci (CAG: n=11, GGC: n=9, AAAAT: n=6)) (Figure 1A, CAG and GGC). This supports our hypothesis that disease-causing tandem repeats are more polymorphic among the normal population. Next we plotted the variation of repeat length (interquartile range (IQR) of repeat-unit count from each read), and mean repeat length, at each exonic locus (including UTR). Shorter-unit repeats are more numerous and more variable (Figure S2). Given that different repeat sequences may have different mutation rates [8], we compared the ten kinds of triplet repeat (Figure S3). Most of the non-disease triplet repeats have little or no length polymorphism. A large fraction (>94% of all repeats) have IQR less than 2, while disease causing tandem repeats usually show more variation (always more than 2) (Figure1B, C). It is of interest that GGC and CAG repeats have more polymorphic loci than other repeat structures (Figure S3).

**Figure1.**
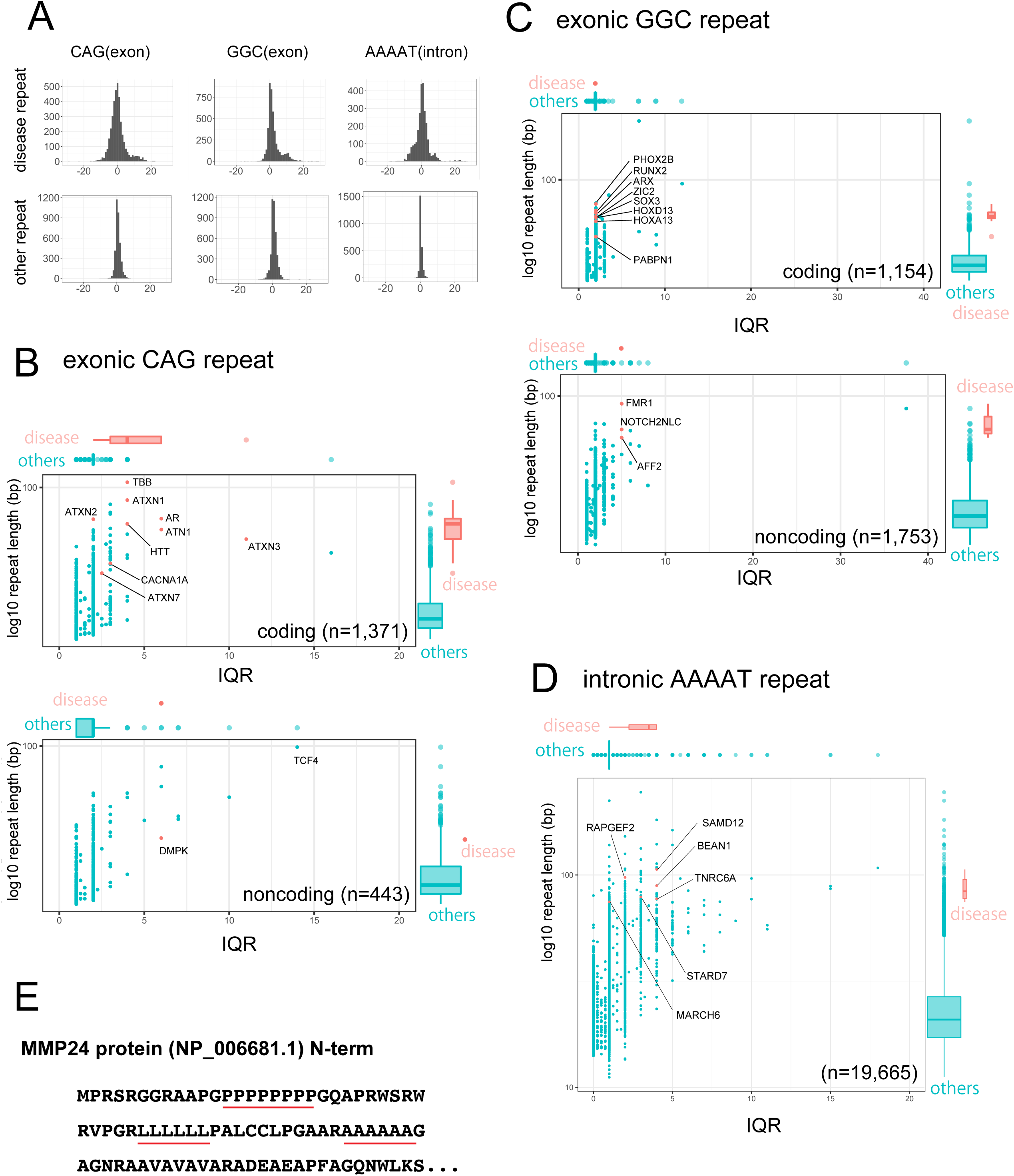
(A) Variation of tandem repeat length (copy number). x-axis: copy number change relative to the human reference (hg38). y-axis: read count. Three different repeat types (exonic CAG, exonic GGC and intronic AAAAT) are separately shown. Disease repeats: 9 coding CAG repeats, 8 coding and 3 5’UTR GGC repeats, and 6 intronic AAAAT repeats. Other repeat: the same number of repeat loci (CAG: n=11, GGC: n=9, AAAAT: n=6) are randomly extracted from all repeat loci (CAG: n=1814, GGC: n=2907, AAAAT: n=19,665) for comparison to the disease repeats (GGC: n=9, CAG: n=11, AAAAT: n=6). (B-D) Variation (IQR) and length of repeats with disease-associated sequences. (B) Upper: coding CAG repeats, Bottom: non-coding exonic CAG repeats. (C) Upper: coding GGC repeats, Bottom: non-coding exonic GGC repeats. (D) Intronic AAAAT repeats. Many of the disease-causing repeats have large variation and long repeat size. x-axis: IQR, y-axis: mean repeat length (bp). n provides the numbers of repeat loci. x-axis: IQR, y-axis: read count. (E) N-terminus amino acid sequence of MMP24. There are poly-leucine, poly-alanine and poly-proline tracts (red lines).

All 9 disease-causing CAG repeats (Table S2) have IQR >= 2 (median: 4, range: 2-11, Figure 1B). Note that all disease causing CAG repeats, except for *DMPK*, are located in protein-coding regions (Figure 1B, Table S2). Importantly, among non-disease CAG repeats, we found a non-coding CAG repeat in *TCF4* with high IQR. This repeat was not included in our rare disease list but it has an association with Fuchs endothelial corneal dystrophy (FECD) (MIM#613267). FECD is a commonly seen disease affecting 4-5% of the population older than 40 years [9]. Initially, genome wide association studies (GWAS) showed an association of a SNP (rs613872), but later studies showed this disease has much higher association to a 4.3kb-downstream CAG repeat [10] [11]. This triplet repeat was known to be highly polymorphic [12], in agreement with our result.

Disease-causing GGC repeats are either in protein-coding or 5’-UTR regions. All known protein-coding GGC repeat diseases are caused by poly-alanine expansions. These poly-alanine loci show less variability (IQR=2) than the 5’-UTR GGC loci (IQR=5) (Figure 1C). This may reflect the difference in disease mechanisms of protein-coding and 5’-UTR GGC repeats. It is known that poly-alanine is highly toxic to cells [13] and usually fewer than 10 additional alanine residues are enough to cause disease [2]. This may explain our observation that alanine-coding GGCs are less variable in the general population. In contrast, disease-causing 5’UTR GGCs are more polymorphic. One possible pathomechanism of 5’UTR GGC repeats is gene suppression as seen in fragile X syndrome [5]. Another envisioned mechanism is repeat associated non-AUG translation, which is suspected in the neurological symptoms in patients with *FMR1* premutation (more than 55 GGC repeats). The different mechanisms may show different variation patterns of disease-causing GGC repeats.

In addition to triplet repeats, pathogenic expansions of quintuplet repeat loci (represented as AAAAT in hg38) are associated with myoclonic epilepsies [14-16]. In 2018 and 2019, five AAAAT repeat loci were reported [14-16] in addition to *BEAN1* which causes spinocerebellar ataxia 31 (MIM#117210) [17]. We also observed that disease-linked AAAAT repeat loci have variation in length (Figure 1A, AAAAT). We examined the variation and length of all intronic AAAAT repeat loci in 16 individuals, and found several highly polymorphic AAAAT repeats including disease loci (IQR=4: *SAMD12, BEAN1, TNRC6A*, Figure 1D). Quintuplet AAAAT repeat expansions are associated with newly-discovered types of disease, and pathomechanisms of AAAAT repeat expansions are yet unclear. It may be that undiscovered pathogenic repeats for epilepsy are among the other highly polymorphic quintuplet repeats.

We repeated our analysis using repeat annotations from Tandem Repeats Finder (TRF, a.k.a. simpleRepeat.txt) [18]. The proportions of triplet repeat sequences were similar to those from `tantan` (Figure S4A). We analyzed disease-causing CAG and GGC repeats, and observed similar results to tantan-annotated repeats (Figure S4B-D), although the *TCF4* repeat was not annotated by TRF (Figure S4B).

GWAS studies have identified numerous genomic markers over the past fifteen years, however their functional relation to the diseases or traits is usually unclear. It is plausible that tandem repeats near those GWAS markers actually have functional relation to the traits. Our approach found one such example, the *TCF4* repeat for corneal disease, so there may be new candidates among the highly polymorphic repeats (Table 1, Figure 1B). We listed highly polymorphic exonic triplet repeats (IQR>=5) near GWAS signals (<10 kb) from a GWAS catalog [19] (Table 1). It is possible that polymorphic tandem repeats contribute to gene expression variation [20, 21]. A recent study showed that tandem repeats which can alter expression of near-by genes are potential drivers of published GWAS signals. Fotsing *et al*. listed such 1380 tandem repeats as eSTR (repeats associated with the expression of nearby genes) [21], although no Mendelian disease-causing repeats are included in eSTR, possibly because until today most of the known repeat diseases are not caused by altering gene expression levels but by changing protein products. However, there may be more diseases or traits caused by altering gene expression, like Fragile X syndrome. We found an interesting candidate, a 5’UTR GCA repeat in the *GLS* gene, which is highly polymorphic and also listed as an eSTR. Several lines of evidence show that an 8kb-downstream SNP is associated with reticulocyte count (Table S3). *GLS* encodes glutaminase, which catalyzes glutamine conversion to glutamate, has high activity in red blood cells, and plays a role in glutathione metabolism [22] [23]. It is an intriguing possibility that this 5’UTR repeat actually acts as a driver of the GWAS signals and affects erythrocyte maturation by altering the expression of *GLS* thus affecting glutathione metabolism. Another candidate, which does not seem to alter gene expression levels but may alter protein function, is *MMP24*. Three CAG or GGC repeats in *MMP24* are highly variable, which has not been reported previously (Table 1). These encode poly-leucine, poly-alanine (neither are annotated by TRF) and poly-proline tracts (Figure 1E, Figure S5). Two SNPs (rs2425019, rs747202389) 4.5kb and 7.5kb downstream of these repeats, respectively, are reported to be associated with height [24]. *MMP24* encodes a membrane matrix metalloprotease and has roles in embryonic development [25]. It would be interesting to investigate functional consequences of changing these repeats. These speculative examples need further association studies targeting near-by tandem repeats together with functional studies to elucidate the mechanistic relation to the phenotype.

**Table 1.**
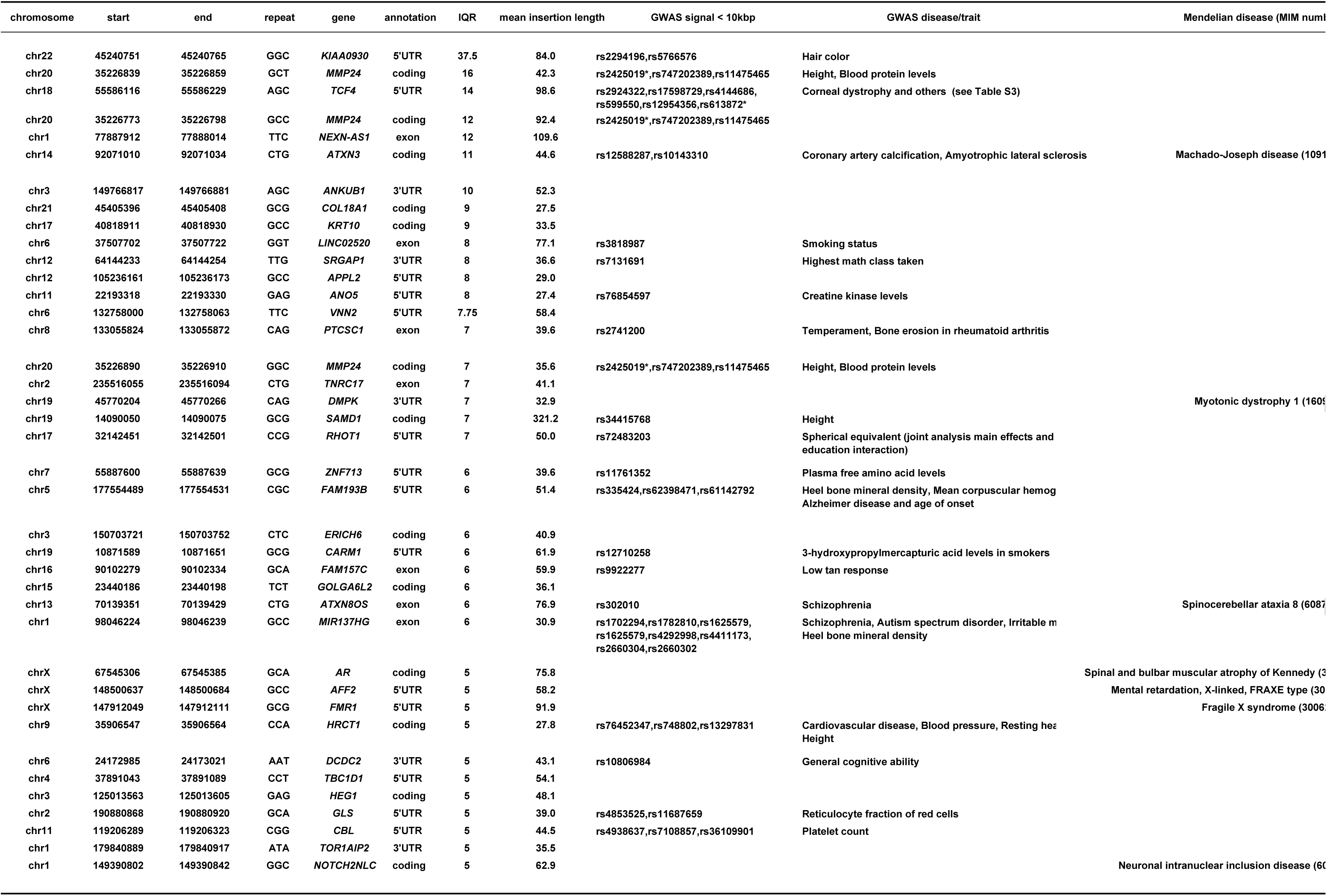
Highly polymorphic triplet repeats (IQR>=5). Repeats with nearby (<10 kb) reported GWAS signals are shown. Repeats whose expansion are known to cause Mendelian disease are also shown. IQR: interquartile range. * not included in GWAS catalog.

In conclusion, our results indicate that the disease-causing coding CAG repeats, 5’UTR GGC repeats, and intronic AAAAT repeats are long and variable, but alanine-coding GGC repeats are stable (but long) among the 16 individuals. In addition, we detected highly polymorphic tandem repeats that are associated with common disease (i.e. *TCF4* repeats). This suggests that polymorphic tandem repeats may often contribute to common diseases. Our study is limited due to lack of a large number of healthy individuals from multiple ethnicities. Nevertheless, we provide a first example of applying long read sequencing to identify polymorphic tandem repeats. We believe further tandem-repeat surveys using a large number of individuals may provide more insights into human genomes and diseases.

## Declarations

### Ethics approval and consent to participate

All genomic DNA were examined after obtaining informed consent. Experimental protocols were approved by institutional review board of Yokohama City University under the number of A19080001.

### Availability of data and materials

The sequence datasets generated and analyzed in this study are not publicly available.

## Author contributions

SM, MCF, and NM contributed to the conception of the work and acquisition/analysis/interpretation of the data.

