## Supplemental materials for "Genome-wide survey of tandem repeats by nanopore sequencing shows that disease-associated repeats are more polymorphic in the general population"

**Supplemental Methods**

**Long read sequencing**

All long read whole genome sequencing datasets of non-disease or non-repeat disease controls were obtained using Nanopore PromethION sequencers as previously described [1], except for two public datasets [2] [3]. All 16 human control whole genome long read sequencing datasets were obtained from healthy or non-repeat disease individuals (e.g. chromosomal translocations).

**tandem repeat prediction**

Tandem repeats in the human reference genome GRCh38 were predicted using `tantan` (http://cbrc3.cbrc.jp/~martin/tantan/), with this command:
tantan -f4 hg38 > tantan-out

**tandem-genotypes**

tandem-repeat copy number changes relative to the reference were predicted using tandem-genotypes and LAST.
tandem-genotypes -g refFlat.txt tantan-out alignnment.maf > out

All `tandem-genotypes` output files from 16 datasets were merged like this:
tandem-genotypes-join file1 file2 file3... > merged-file

IQR and mean length were calculated from `tandem-genotypes` output using GNU datamash (https://www.gnu.org/software/datamash/).

#### Figure legends

**Supplemental Figure 1. Detection rate of all the tandem repeats.**

(A) Number of `tantan`-annotated tandem repeats for each repeat unit length. y-axis: log10 number of repeat loci.

(B) Detection rates in each repeat unit category are shown. Detection rate is the percentage of repeats whose length is predicted from at least one read.

**Supplemental Figure 2. Variability of exonic repeats.**

Number of exonic tandem repeats (A) and IQR (B) of each unit. Shorter-unit repeats have more variation. Dots represent outliers. Boxplot ranges are the 25th and 75th percentiles. Lines in boxes are median. Most of the IQRs from repeats whose length are more than six are zero.

**Supplemental Figure 3. Deviation and repeat length of each triplet repeat type.**

There are 10 kinds of triplet repeat, AAC, CAC, CCT, CTT, GAT, GTA, GTC, TAA CAG and GGC. The number of repeats are; 580 (AAC), 615 (CAC), 1833 (CCT), 711 (CTT), 417 (GAT), 43 (GTA), 72 (GTC), 575 (TAA), 1814 (CAG) and 2907 (GGC). The variation of the repeat length in 16 individuals were separately shown. x-axis: IQR, y-axis: mean repeat length (bp). In merged boxplots, ranges are the 25th and 75th percentiles, dots are outliers and lines in boxes are median.

**Supplemental Figure 4. Repeat variability using TRF-annotated repeats.**

(A) Proportions of triplet repeat sequences are similar between `tantan` and TRF annotation. (B-D) Variation and length of repeats with disease-associated sequences. For `tandem-genotypes` analysis, we used TRF annotation. Exonic CAG (B), exonic GGC (C) and intronic AAAAT (D) repeats. Many of the disease-causing repeats have large variation and long repeat size. x-axis: IQR, y-axis: repeat length (bp). x-axis: IQR, y-axis: read count.

**Supplemental Figure 5. MMP24 has three polymorphic protein-coding repeats.**

MMP24 has three protein coding repeats near the N-terminus of the protein. These are poly-proline, poly-leucine and poly-alanine tracts. The lower pictures show them in the UCSC genome browser with comparison to other mammals. They are completely conserved in the chimpanzee genome, and partly conserved in mouse.

Figure S1

A

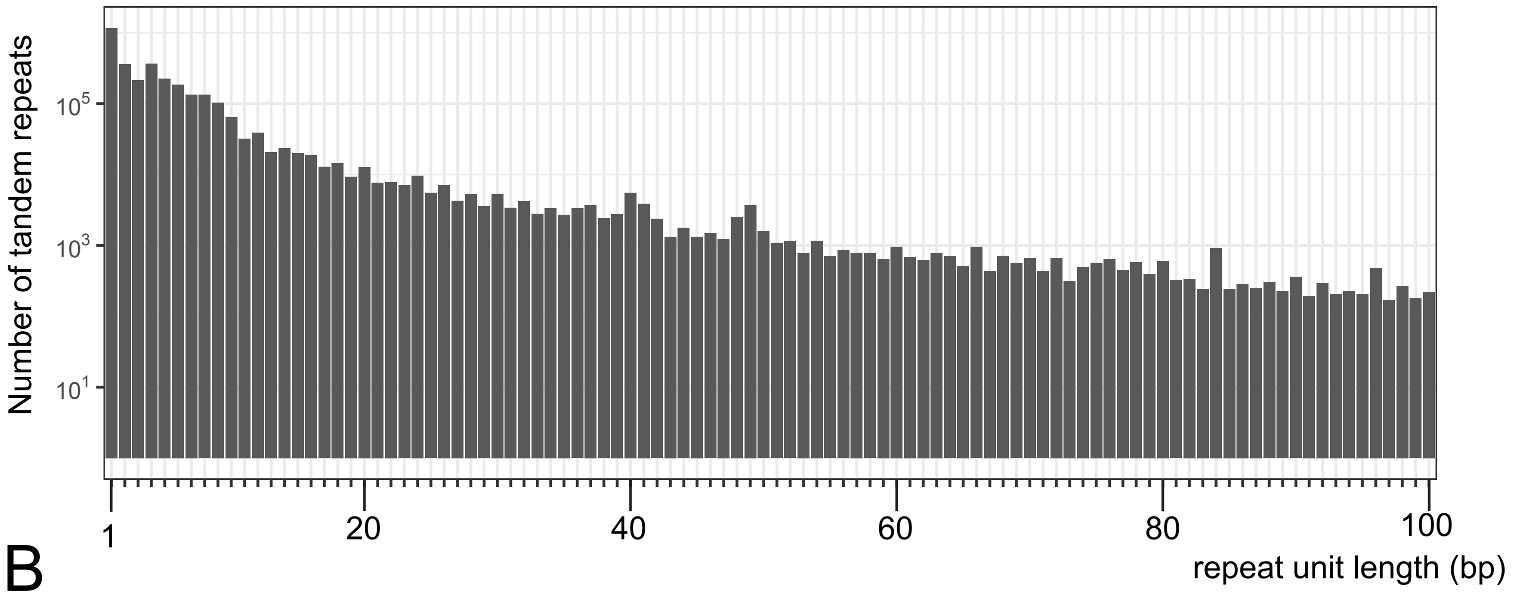

Number of tandem repeats
$10^5$
$10^3$
$10^1$
1
20
40
60
80
100
repeat unit length (bp)

B

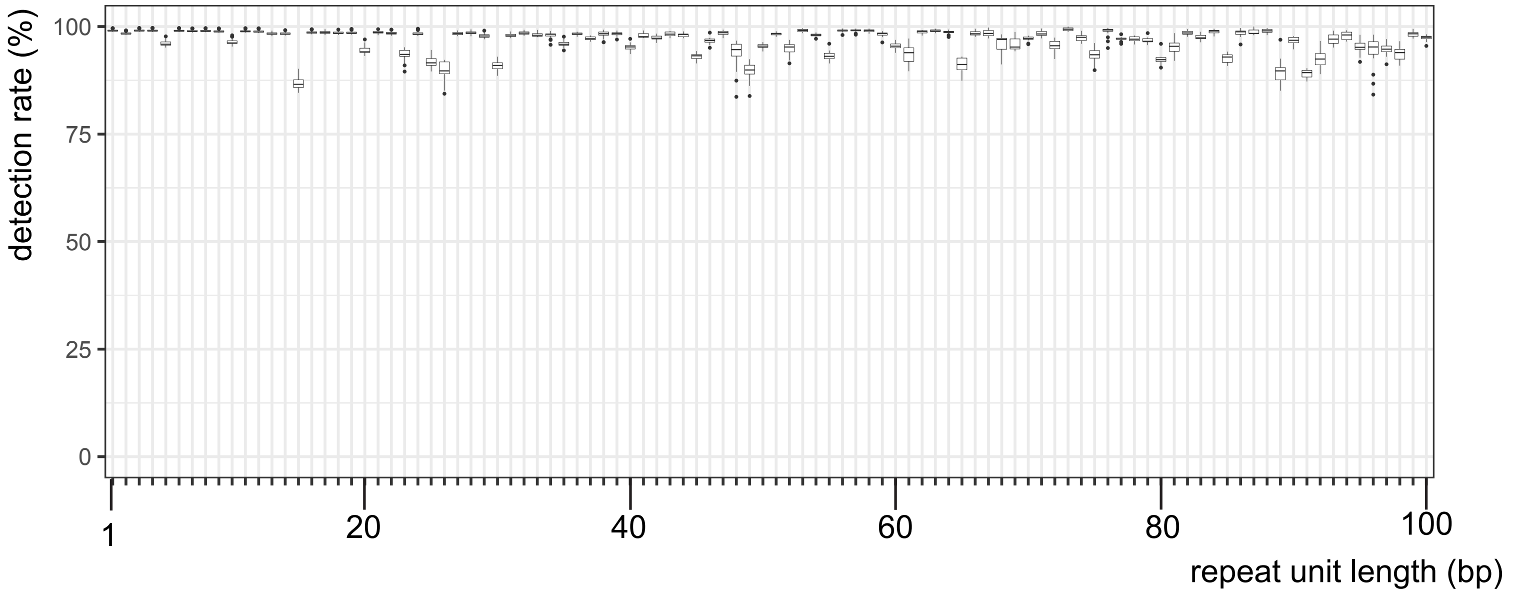

detection rate (%)
100
75
50
25
0
1
20
40
60
80
100
repeat unit length (bp)

### Figure S2

A

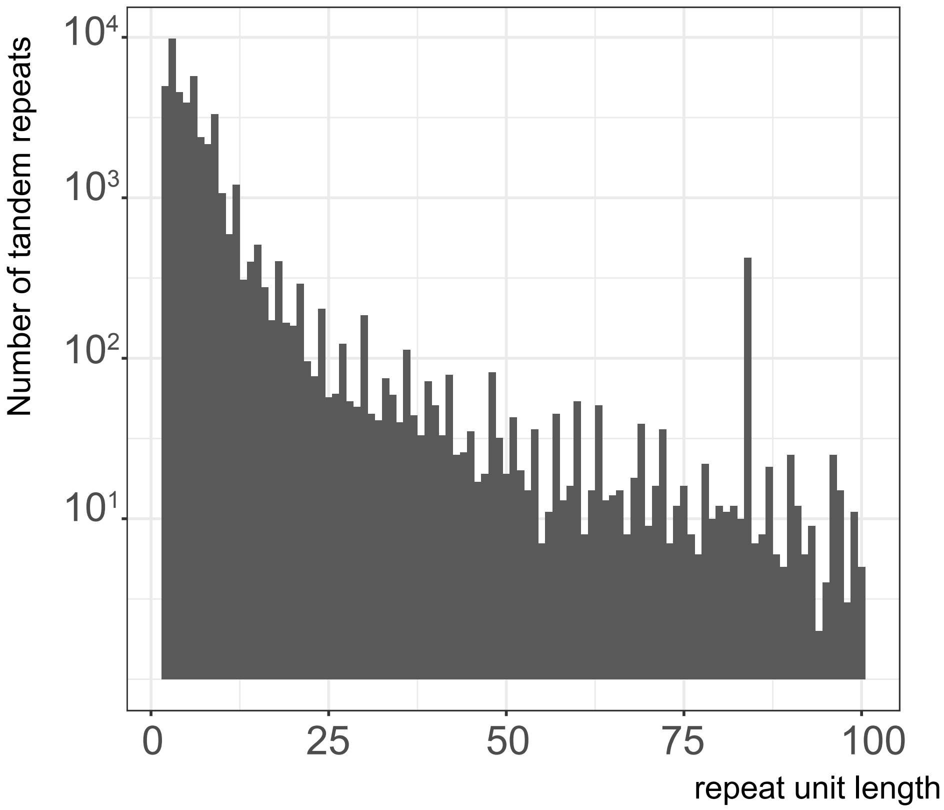

10⁴
10³
10²
10¹
Number of tandem repeats
0
25
50
75
100
repeat unit length

B

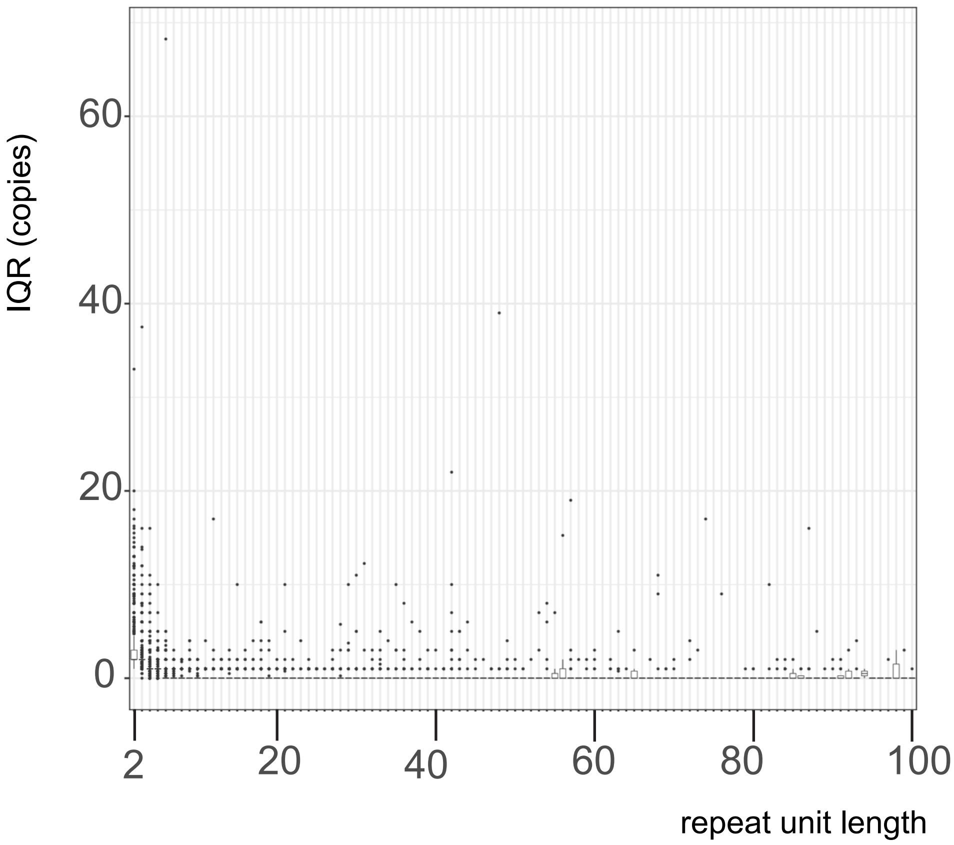

60
40
20
0
IQR (copies)
2
20
40
60
80
100
repeat unit length

### Figure S3

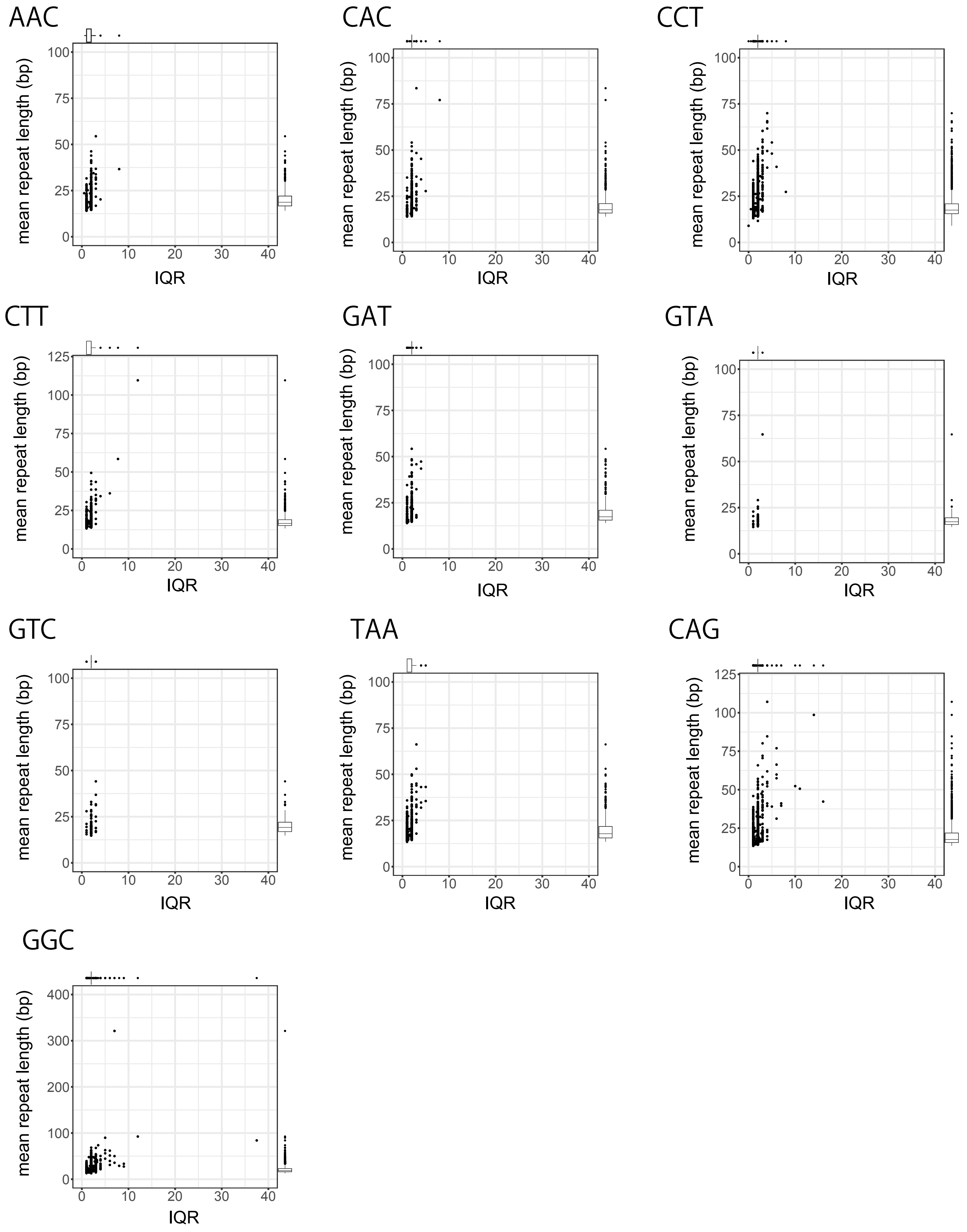

AAC
CAC
CCT
CTT
GAT
GTA
GTC
TAA
CAG
GGC
mean repeat length (bp)
IQR

### Figure S4

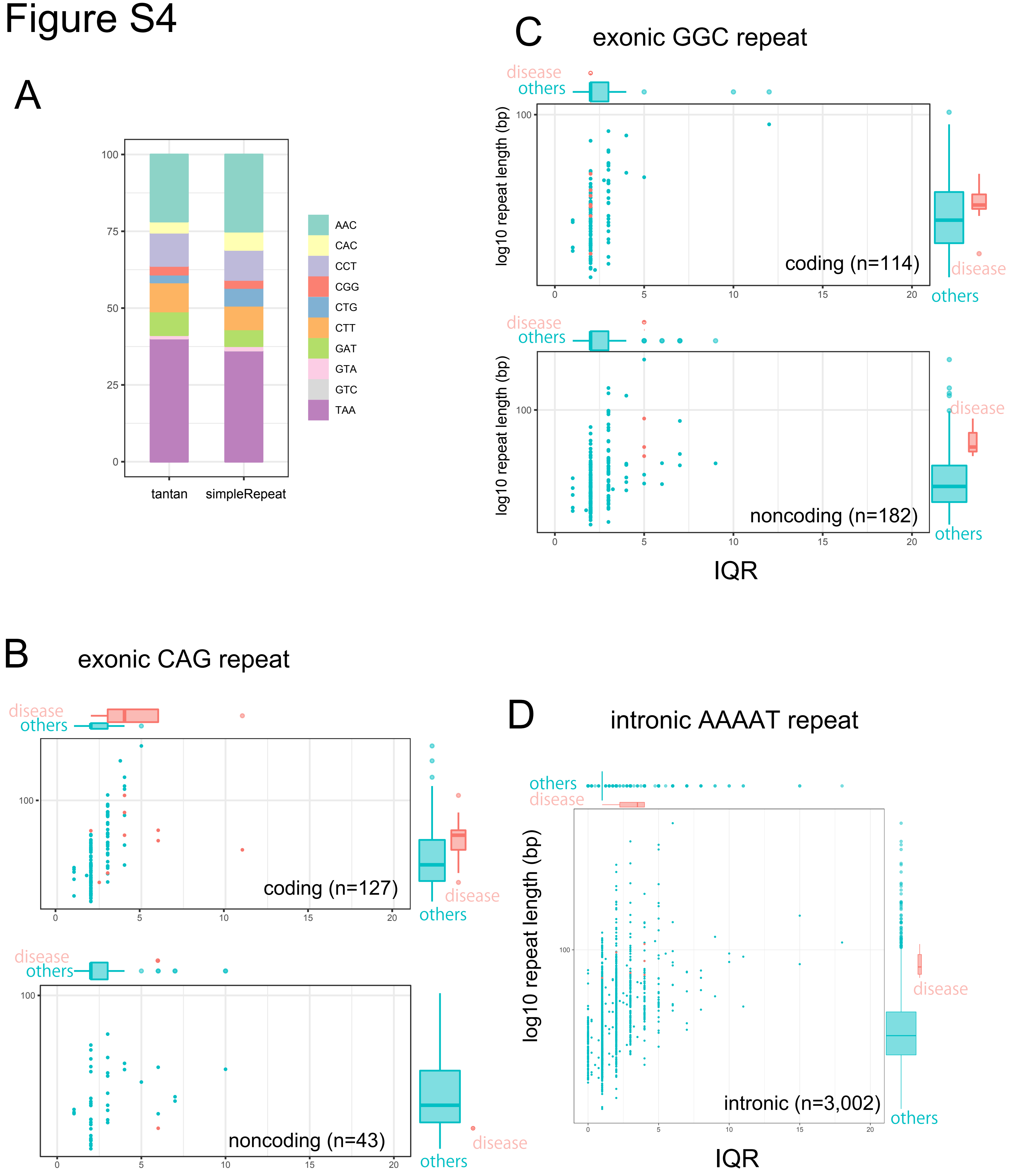
A
100
75
50
25
0
tantan
simpleRepeat
AAC
CAC
CCT
CGG
CTG
CTT
GAT
GTA
GTC
TAA
C
exonic GGC repeat
disease
others
log10 repeat length (bp)
100
coding (n=114)
0
5
10
15
20
disease
others
disease
others
log10 repeat length (bp)
100
noncoding (n=182)
0
5
10
15
20
disease
others
IQR
B
exonic CAG repeat
disease
others
100
coding (n=127)
0
5
10
15
20
disease
others
disease
others
100
noncoding (n=43)
0
5
10
15
20
disease
others
D
intronic AAAAT repeat
others
disease
log10 repeat length (bp)
100
intronic (n=3,002)
0
5
10
15
20
disease
others
IQR

### Figure S5

##### MMP24 protein (NP_006681.1)

**MPRSRGGRAAPGPPPPPPPPGQAPRWSRWRVPGRLLLLLLPALCCLPGAARAAAAAA**

**GAGNRAAVAVAVARADEAEAPFAGQNWLKSYGYLLPYDSRASALHSAKALQSAVSTM**

**QQFYGIPVTGVLDQTTIEWMKKPRCGVPDHPHLSRRRRNKRYALTGQKWRQKHITYS**

**IHNYTPKVGELDTRKAIRQAFDVWQKVTPLTFEEVPYHEIKSDRKEADIMIFFASGF**

**HGDSSPFDGEGGFLAHAYFPGPGIGGDTHFDSDEPWTLGNANHDGNDLFLVAVHELG**

**HALGLEHSSDPSAIMAPFYQYMETHNFKLPQDDLQGIQKIYGPPAEPLEPTRPLPTL**

**PVRRIHSPSERKHERQPRPPRPPLGDRPSTPGTKPNICDGNFNTVALFRGEMFVFKD**

**RWFWRLRNNRVQEGYPMQIEQFWKGLPARIDAAYERADGRFVFFKGDKYWVFKEVTV**

**EPGYPHSLGELGSCLPREGIDTALRWEPVGKTYFFKGERYWRYSEERRATDPGYPKP**

**ITVWKGIPQAPQGAFISKEGYYTYFYKGRDYWKFDNQKLSVEPGYPRNILRDWMGCN**

**QKEVERRKERRLPQDDVDIMVTINDVPGSVNAVAVVIPCILSLCILVLVYTIFQFKN**

**KTGPQPVTYYKRPVQEWV**

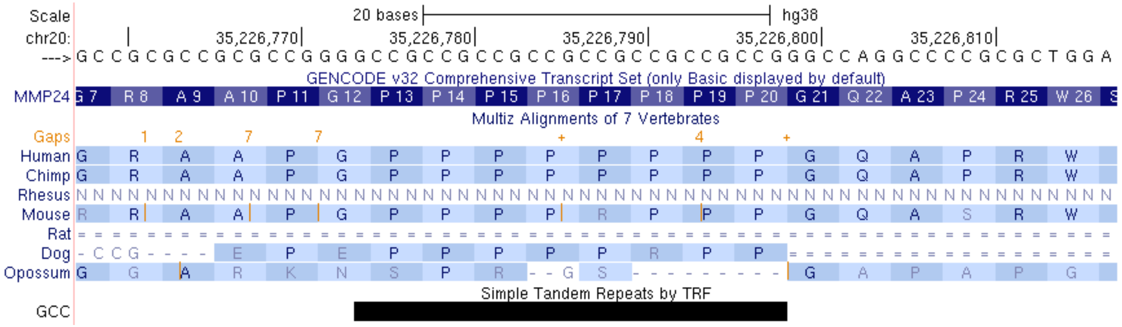

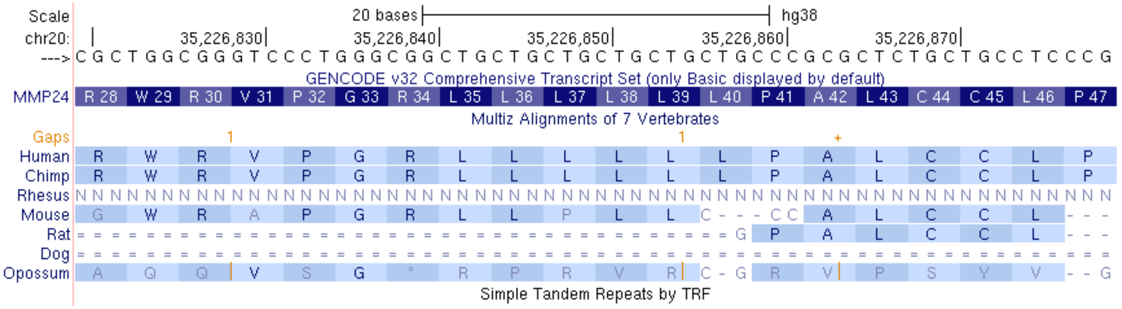

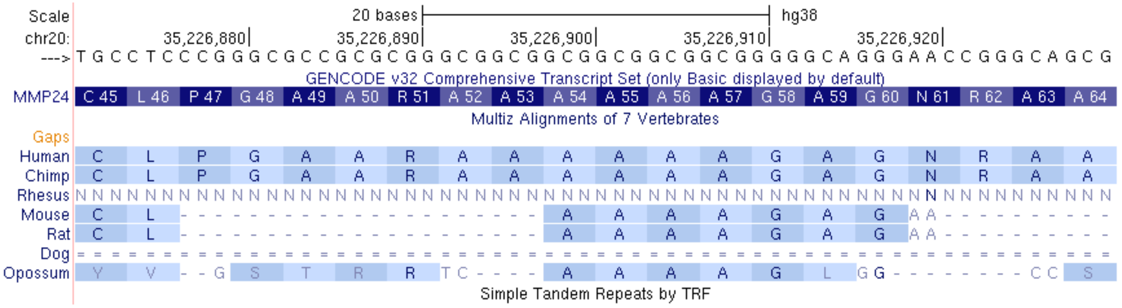

| data # | | | Ethnicity | data size (Gb) | detection rate of all tandem-repeat*** |
|---|---|---|---|---|---|
| data1* | rel3 | MinION | Caucasian | 91 | 98.4 |
| data2** | ERR258112-5 | PromethION | Yorban | 185 | 98.5 |
| data3 | prom5 | PromethION | Japanese | 90 | 99.2 |
| data4 | prom6 | PromethION | Japanese | 81 | 98.4 |
| data5 | prom7 | PromethION | Japanese | 111 | 98.4 |
| data6 | prom9 | PromethION | Japanese | 35 | 98.0 |
| data7 | prom11 | PromethION | Japanese | 86 | 99.1 |
| data8 | prom12 | PromethION | Japanese | 54 | 98.2 |
| data9 | prom14 | PromethION | Japanese | 53 | 99.1 |
| data10 | prom24 | PromethION | Japanese | 63 | 98.2 |
| data11 | prom31 | PromethION | Japanese | 41 | 98.3 |
| data12 | prom32 | PromethION | Japanese | 94 | 98.4 |
| data13 | prom33 | PromethION | Japanese | 117 | 98.4 |
| data14 | prom37 | PromethION | Chinese | 75 | 99.2 |
| data15 | prom65 | PromethION | Japanese | 55 | 98.4 |
| data16 | prom66 | PromethION | Japanese | 72 | 98.4 |

**Table S1**. Data sets used in this study. Detection rates are the number of tandem repeats whose length is predicted from at least one DNA read. Data 1 and 2 are public nanopore data. data1*: rel3 [3], data2**: ERR258112-5 [2], *** 3,312,291 loci.

| chromosome | start | end | repeat | gene | function | Disease | OMIM | IQR (copies) | mean length (bp) |
|---|---|---|---|---|---|---|---|---|---|
| chr2 | 176093058 | 176093103 | GGC | *HOXD13* | coding | Syndactyly, type V | 186300 | 2 | 47.3 |
| chr4 | 41745971 | 41746031 | GGC | *PHOX2B* | coding | Congenital central hypoventilation syndrome | 209880 | 2 | 62.5 |
| chr14 | 23321472 | 23321502 | GGC | *PABPN1* | coding | Oculopharyngeal muscular dystrophy | 164300 | 2 | 32.8 |
| chr6 | 45422750 | 45422801 | GGC | *RUNX2* | coding | Cleidocranial dysplasia, forme fruste, with brachydactyly | 119600 | 2 | 54.1 |
| chrX | 25013649 | 25013697 | GGC | *ARX* | coding | Early infantile epileptic encephalopathy | 308350 | 2 | 51.6 |
| chr13 | 99985448 | 99985493 | GGC | *ZIC2* | coding | Holoprosencephaly | 609637 | 2 | 48.0 |
| chrX | 140504316 | 140504361 | GGC | *SOX3* | coding | Mental retardation with isolated growth hormone deficiency | 300123 | 2 | 47.8 |
| chr7 | 27199924 | 27199966 | GGC | *HOXA13* | coding | Hand-foot-genital syndrome | 140000 | 2 | 43.8 |
| chr1 | 149390802 | 149390842 | GGC | *NOTCH2NLC* | 5'UTR | Neuronal intranuclear inclusion disease | 603472 | 5 | 62.9 |
| chrX | 147912050 | 147912110 | GGC | *FMR1* | 5'UTR | Fragile X syndrome/tremor-ataxia syndrome | 300624/300623 | 5 | 89.9 |
| chrX | 148500637 | 148500682 | GGC | *AFF2* | 5'UTR | Fragile X syndrome | 309548 | 5 | 55.9 |
| chr14 | 92071010 | 92071040 | CAG | *ATXN3* | coding | Spinocerebellar ataxia 3 | 109150 | 11 | 50.5 |
| chr19 | 13207858 | 13207897 | CAG | *CACNA1A* | coding | Spinocerebellar ataxia 6 | 183086 | 3 | 36.7 |
| chr12 | 6936716 | 6936773 | CAG | *ATN1* | coding | Dentatorubral-pallidoluysian atrophy | 125370 | 6 | 56.9 |
| chr6 | 16327635 | 16327722 | CAG | *ATXN1* | coding | Spinocerebellar ataxia 1 | 164400 | 4 | 84.4 |
| chr3 | 63912685 | 63912715 | CAG | *ATXN7* | coding | Spinocerebellar ataxia 7 | 164500 | 3 | 32.2 |
| chr6 | 170561907 | 170562021 | CAG | *TBP* | coding | Spinocerebellar ataxia 17 | 607136 | 4 | 106.8 |
| chr12 | 111598950 | 111599019 | CAG | *ATXN2* | coding | Spinocerebellar ataxia 2 | 183090 | 2 | 65.8 |
| chr4 | 3074876 | 3074939 | CAG | *HTT* | coding | Huntington disease | 143100 | 4 | 61.6 |
| chrX | 67545317 | 67545386 | CAG | *AR* | coding | Spinal and Bulbar Muscular Atrophy | 313200 | 5.25 | 66.1 |
| chr16 | 66490396 | 66490466 | AAAAT | *BEAN1* | intron | Spinocerebellar ataxia 31 | 117210 | 4 | 88.3 |
| chr8 | 118366815 | 118366918 | AAAAT | *SAMD12* | intron | Epilepsy, familial adult myoclonic, 1 | 601068 | 4 | 105.3 |
| chr4 | 159342526 | 159342618 | AAAAT | *RAPGEF2* | intron | Epilepsy, familial adult myoclonic, 7 | 618075 | 2 | 96.5 |
| chr16 | 24613438 | 24613532 | AAAAT | *TNRC6A* | intron | Epilepsy, familial adult myoclonic, 6 | 618074 | 4 | 76.8 |
| chr5 | 10356339 | 10356411 | AAAAT | *MARCH6* | intron | Myoclonic epilepsy | - | 1 | 74.3 |
| chr2 | 96197066 | 96197124 | AAAAT | *STARD7* | intron | Myoclonic epilepsy | - | 3 | 78.5 |

**Table S2**. Variability of triplet- and quintuplet-repeat disease locus in 16 individuals. OMIM: Online Mendelian Inheritance in Men.

| | | | | | | | | | GWAS-catalog | | | | | | | | | |
|---|---|---|---|---|---|---|---|---|---|---|---|---|---|---|---|---|---|---|
| hromosome | start | end | repeat | gene | context | IQR (copies) | mean insertion length (bp) | Distance from the repeat (bp) | DISEASE/TRAIT | CHR_ID | CHR_POS | SNPS | RISK ALLELE FREQUENCY | P-VALUE | PVALUE_MLOG | OR or BETA | PUBMEDID | FIRST AUTH |
| chr22 | 45240751 | 45240765 | GGC | KIAA0930 | 5'UTR | 37.5 | 84.0 | -5970 | Blond vs. brown/black hair color | 22 | 45234781 | rs2294196 | NR | 2.00E-09 | 8.698970004 | 1.0469012 | 30531825 | Morgan MD |
| chr22 | 45240751 | 45240765 | GGC | KIAA0930 | 5'UTR | 37.5 | 84.0 | -5239 | Hair color | 22 | 45235512 | rs5766576 | NR | 3.00E-08 | 7.522878745 | | 30595370 | Kichaev G |
| chr20 | 35226839 | 35226859 | GCT | MMP24 | coding | 16 | 42.3 | -7524 | Height | 20 | 35219315 | rs747202389 | 0.8057 | 3.00E-06 | 5.522878745 | 0.0499 | 28552196 | Tachmazido |
| chr20 | 35226839 | 35226859 | GCT | MMP24 | coding | 16 | 42.3 | -3832 | Blood protein levels | 20 | 35223007 | rs11475465 | 0.96 | 2.00E-56 | 55.69897 | 1.15 | 29875488 | Sun BB |
| chr18 | 55586116 | 55586229 | AGC | TCF4 | 5'UTR | 14 | 98.6 | -8933 | Hand grip strength | 18 | 55577183 | rs2924322 | 0.119 | 6.00E-08 | 7.22184875 | 0.1859 | 29313844 | Willems SM |
| chr18 | 55586116 | 55586229 | AGC | TCF4 | 5'UTR | 14 | 98.6 | -8933 | Self-rated health | 18 | 55577183 | rs2924322 | NR | 5.00E-08 | 7.301029996 | 0.039 | 27864402 | Harris SE |
| chr18 | 55586116 | 55586229 | AGC | TCF4 | 5'UTR | 14 | 98.6 | -1785 | Schizophrenia | 18 | 55584331 | rs17598729 | NR | 9.00E-11 | 10.04575749 | 1.0893246 | 30285260 | Ikeda M |
| chr18 | 55586116 | 55586229 | AGC | TCF4 | 5'UTR | 14 | 98.6 | -1622 | Smoking behaviour (cigarettes smoked per day) | 18 | 55584494 | rs4144686 | 0.167 | 1.00E-08 | 8 | 0.018551348 | 30643251 | Liu M |
| chr18 | 55586116 | 55586229 | AGC | TCF4 | 5'UTR | 14 | 98.6 | -1622 | Cigarettes smoked per day (MTAG) | 18 | 55584494 | rs4144686 | 0.167 | 9.00E-09 | 8.045757491 | 0.0168007 | 30643251 | Liu M |
| chr18 | 55586116 | 55586229 | AGC | TCF4 | 5'UTR | 14 | 98.6 | -959 | Feeling fed-up | 18 | 55585157 | rs599550 | 0.147785 | 3.00E-14 | 13.52287875 | 7.6 | 29500382 | Nagel M |
| chr18 | 55586116 | 55586229 | AGC | TCF4 | 5'UTR | 14 | 98.6 | -959 | Feeling lonely | 18 | 55585157 | rs599550 | 0.147785 | 7.00E-11 | 10.15490196 | 6.53 | 29500382 | Nagel M |
| chr18 | 55586116 | 55586229 | AGC | TCF4 | 5'UTR | 14 | 98.6 | -959 | Pulse pressure | 18 | 55585157 | rs599550 | 0.859 | 7.00E-10 | 9.15490196 | 0.1747 | 30578418 | Giri A |
| chr18 | 55586116 | 55586229 | AGC | TCF4 | 5'UTR | 14 | 98.6 | -959 | Depressed affect | 18 | 55585157 | rs599550 | NR | 4.00E-17 | 16.39794001 | 0.02725 | 29942085 | Nagel M |
| chr18 | 55586116 | 55586229 | AGC | TCF4 | 5'UTR | 14 | 98.6 | -959 | Systolic blood pressure | 18 | 55585157 | rs599550 | 0.8602 | 5.00E-13 | 12.30103 | 0.2834 | 30578418 | Giri A |
| chr18 | 55586116 | 55586229 | AGC | TCF4 | 5'UTR | 14 | 98.6 | 63 | Autism spectrum disorder or schizophrenia | 18 | 55586179 | rs12954356 | | 9.00E-10 | 9.045757491 | 1.0869565 | 28540026 | Anney RJL |
| chr20 | 35226773 | 35226798 | GCC | MMP24 | coding | 12 | 92.4 | -7458 | Height | 20 | 35219315 | rs747202389 | 0.8057 | 3.00E-06 | 5.522878745 | 0.0499 | 28552196 | Tachmazido |
| chr20 | 35226773 | 35226798 | GCC | MMP24 | coding | 12 | 92.4 | -3766 | Blood protein levels | 20 | 35223007 | rs11475465 | 0.96 | 2.00E-56 | 55.69897 | 1.15 | 29875488 | Sun BB |
| chr14 | 92071010 | 92071034 | CTG | ATXN3 | coding | 11 | 44.6 | -395 | Coronary artery calcification | 14 | 92070615 | rs12588287 | 0.83 | 9.00E-06 | 5.045757491 | 0.19 | 23870195 | Wojczynski |
| chr14 | 92071010 | 92071034 | CTG | ATXN3 | coding | 11 | 44.6 | 3027 | Amyotrophic lateral sclerosis | 14 | 92074037 | rs10143310 | 0.2436 | 3.00E-07 | 6.522878745 | 1.09 | 29566793 | Nicolas A |
| chr6 | 37507702 | 37507722 | GGT | LINC02520 | exon | 8 | 77.1 | 9128 | Smoking status | 6 | 37516830 | rs3818987 | NR | 6.00E-10 | 9.22184875 | | 30595370 | Kichaev G |
| chr6 | 37507702 | 37507722 | GGT | LINC02520 | exon | 8 | 77.1 | 9128 | Smoking status (ever vs never smokers) | 6 | 37516830 | rs3818987 | 0.4734 | 3.00E-10 | 9.522878745 | 0.013703931 | 30643258 | Karlsson Lir |
| chr12 | 64144233 | 64144254 | TTG | SRGAP1 | 3'UTR | 8 | 36.6 | 2947 | Highest math class taken (MTAG) | 12 | 64147180 | rs7131691 | 0.4702 | 2.00E-08 | 7.698970004 | 0.0093 | 30038396 | Lee JJ |
| chr11 | 22193318 | 22193330 | GAG | ANO5 | 5'UTR | 8 | 27.4 | 6920 | Creatine kinase levels | 11 | 22200238 | rs76854597 | NR | 1.00E-21 | 21 | 0.05561 | 29403010 | Kanai M |
| chr8 | 133055824 | 133055872 | CAG | PTCSC1 | exon | 7 | 39.6 | 3764 | Temperament | 8 | 133059588 | rs2741200 | 0.671 | 5.00E-06 | 5.301029996 | 0.32 | 22832960 | Service SK |
| chr8 | 133055824 | 133055872 | CAG | PTCSC1 | exon | 7 | 39.6 | 3764 | Bone erosion in rheumatoid arthritis | 8 | 133059588 | rs2741200 | NR | 4.00E-06 | 5.397940009 | 2.0833335 | 28512992 | Joo YB |
| chr20 | 35226890 | 35226910 | GGC | MMP24 | coding | 7 | 35.6 | -7575 | Height | 20 | 35219315 | rs747202389 | 0.8057 | 3.00E-06 | 5.522878745 | 0.0499 | 28552196 | Tachmazido |
| chr20 | 35226890 | 35226910 | GGC | MMP24 | coding | 7 | 35.6 | -3883 | Blood protein levels | 20 | 35223007 | rs11475465 | 0.96 | 2.00E-56 | 55.69897 | 1.15 | 29875488 | Sun BB |
| chr19 | 14090050 | 14090075 | GCG | SAMD1 | coding | 7 | 321.2 | 7920 | Height | 19 | 14097970 | rs34415768 | NR | 3.00E-08 | 7.522878745 | | 30595370 | Kichaev G |
| chr17 | 32142451 | 32142501 | CCG | RHOT1 | 5'UTR | 7 | 50.0 | -5585 | Spherical equivalent (joint analysis main effects and education interaction) | 17 | 32136866 | rs72483203 | 0.09 | 9.00E-09 | 8.045757491 | | 27020472 | Fan Q |
| chr17 | 32142451 | 32142501 | CCG | RHOT1 | 5'UTR | 7 | 50.0 | -5585 | Spherical equivalent (joint analysis main effects and education interaction) | 17 | 32136866 | rs72483203 | 0.06 | 5.00E-10 | 9.301029996 | | 27020472 | Fan Q |
| chr17 | 32142451 | 32142501 | CCG | RHOT1 | 5'UTR | 7 | 50.0 | -5585 | Spherical equivalent (joint analysis main effects and education interaction) | 17 | 32136866 | rs72483203 | 0.17 | 9.00E-06 | 5.045757491 | | 27020472 | Fan Q |
| chr7 | 55887600 | 55887639 | GCG | ZNF713 | 5'UTR | 6 | 39.6 | -4346 | Plasma free amino acid levels | 7 | 55883254 | rs11761352 | 0.38 | 9.00E-17 | 16.04575749 | 0.3 | 30659259 | Imaizumi A |
| chr7 | 55887600 | 55887639 | GCG | ZNF713 | 5'UTR | 6 | 39.6 | -4346 | Plasma free amino acid levels (adjusted for twenty other PFAAs) | 7 | 55883254 | rs11761352 | | 4.00E-19 | 18.39794001 | 0.23 | 30659259 | Imaizumi A |
| chr5 | 177554489 | 177554531 | CGC | FAM193B | 5'UTR | 6 | 51.4 | -6350 | Heel bone mineral density | 5 | 177548139 | rs335424 | NR | 3.00E-10 | 9.522878745 | 0.0135462 | 30048462 | Kim SK |
| chr5 | 177554489 | 177554531 | CGC | FAM193B | 5'UTR | 6 | 51.4 | -6350 | Heel bone mineral density | 5 | 177548139 | rs335424 | NR | 4.00E-11 | 10.39794001 | | 30595370 | Kichaev G |
| chr5 | 177554489 | 177554531 | CGC | FAM193B | 5'UTR | 6 | 51.4 | -586 | Mean corpuscular hemoglobin | 5 | 177553903 | rs62398471 | NR | 5.00E-10 | 9.301029996 | | 30595370 | Kichaev G |
| chr5 | 177554489 | 177554531 | CGC | FAM193B | 5'UTR | 6 | 51.4 | 1973 | Alzheimer disease and age of onset | 5 | 177556462 | rs61142792 | NR | 4.00E-07 | 6.397940009 | 5.083 | 26830138 | Herold C |
| chr19 | 10871589 | 10871651 | GCG | CARM1 | 5'UTR | 6 | 61.9 | 8064 | 3-hydroxypropylmercapturic acid levels in smokers | 19 | 10879653 | rs12710258 | NR | 6.00E-07 | 6.22184875 | 0.1876 | 26053186 | Park SL |
| chr16 | 90102279 | 90102334 | GCA | FAM157C | exon | 6 | 59.9 | -9849 | Low tan response | 16 | 90092430 | rs9922277 | NR | 4.00E-139 | 138.39794 | 0.223 | 29739929 | Visconti A |
| chr13 | 70139351 | 70139429 | CTG | ATXN8OS | exon | 6 | 76.9 | 5979 | Schizophrenia | 13 | 70145330 | rs302010 | NR | 2.00E-06 | 5.698970004 | 1.0638298 | 26198764 | Goes FS |
| chr1 | 98046224 | 98046239 | GCC | MIR137HG | exon | 6 | 30.9 | -9796 | Schizophrenia | 1 | 98036428 | rs1702294 | 0.809 | 3.00E-19 | 18.52287875 | 1.1273957 | 25056061 | Ripke S |
| chr1 | 98046224 | 98046239 | GCC | MIR137HG | exon | 6 | 30.9 | -9796 | Schizophrenia | 1 | 98036428 | rs1702294 | NR | 1.00E-17 | 17 | 1.126 | 30285260 | Ikeda M |
| chr1 | 98046224 | 98046239 | GCC | MIR137HG | exon | 6 | 30.9 | -9796 | Schizophrenia | 1 | 98036428 | rs1702294 | NR | 1.00E-17 | 17 | | 31268507 | Periyasamy |
| chr1 | 98046224 | 98046239 | GCC | MIR137HG | exon | 6 | 30.9 | -9440 | Autism spectrum disorder or schizophrenia | 1 | 98036784 | rs1782810 | | 1.00E-19 | 19 | 1.12 | 28540026 | Anney RJL |
| chr1 | 98046224 | 98046239 | GCC | MIR137HG | exon | 6 | 30.9 | -8846 | Schizophrenia | 1 | 98037378 | rs1625579 | 0.8 | 2.00E-11 | 10.69897 | 1.12 | 21926974 | Ripke S |
| chr1 | 98046224 | 98046239 | GCC | MIR137HG | exon | 6 | 30.9 | -8846 | Autism spectrum disorder, attention deficit-hyperactivity disorder, bipolar disorder | 1 | 98037378 | rs1625579 | 0.801 | 2.00E-11 | 10.69897 | | 23453885 | Smoller JW |
| chr1 | 98046224 | 98046239 | GCC | MIR137HG | exon | 6 | 30.9 | -4084 | Heel bone mineral density | 1 | 98042140 | rs4292998 | NR | 3.00E-11 | 10.52287875 | | 30595370 | Kichaev G |
| chr1 | 98046224 | 98046239 | GCC | MIR137HG | exon | 6 | 30.9 | -4062 | Irritable mood | 1 | 98042162 | rs4411173 | 0.162507 | 5.00E-09 | 8.301029996 | 5.85 | 29500382 | Nagel M |
| chr1 | 98046224 | 98046239 | GCC | MIR137HG | exon | 6 | 30.9 | 347 | Schizophrenia | 1 | 98046571 | rs2660304 | NR | 1.00E-17 | 17 | 1.12 | 26198764 | Goes FS |
| chr1 | 98046224 | 98046239 | GCC | MIR137HG | exon | 6 | 30.9 | 347 | Schizophrenia | 1 | 98046571 | rs2660304 | NR | 2.00E-18 | 17.69897 | 1.1148272 | 29483656 | Pardinas AF |
| chr1 | 98046224 | 98046239 | GCC | MIR137HG | exon | 6 | 30.9 | 8439 | Sleep duration | 1 | 98054663 | rs2660302 | 0.187959 | 5.00E-10 | 9.301029996 | 0.03636245 | 30531941 | Doherty A |
| chr9 | 35906547 | 35906564 | CCA | HRCT1 | coding | 5 | 27.8 | -73 | Cardiovascular disease | 9 | 35906474 | rs76452347 | NR | 5.00E-14 | 13.30103 | | 30595370 | Kichaev G |
| chr9 | 35906547 | 35906564 | CCA | HRCT1 | coding | 5 | 27.8 | -73 | Systolic blood pressure | 9 | 35906474 | rs76452347 | NR | 1.00E-11 | 11 | | 30595370 | Kichaev G |
| chr9 | 35906547 | 35906564 | CCA | HRCT1 | coding | 5 | 27.8 | -73 | Diastolic blood pressure | 9 | 35906474 | rs76452347 | 0.191 | 7.00E-10 | 9.15490196 | 0.23 | 27618448 | Liu C |
| chr9 | 35906547 | 35906564 | CCA | HRCT1 | coding | 5 | 27.8 | 2974 | Resting heart rate | 9 | 35909521 | rs748802 | 0.34 | 1.00E-08 | 8 | 0.2022 | 27798624 | Eppinga RN |
| chr9 | 35906547 | 35906564 | CCA | HRCT1 | coding | 5 | 27.8 | 5289 | Height | 9 | 35911836 | rs13297831 | NR | 5.00E-29 | 28.30103 | | 30595370 | Kichaev G |
| chr6 | 24172985 | 24173021 | AAT | DCDC2 | 3'UTR | 5 | 43.1 | 8618 | General cognitive ability | 6 | 24181603 | rs10806984 | NR | 3.00E-10 | 9.522878745 | 6.284 | 29844566 | Davies G |
| chr2 | 190880868 | 190880920 | GCA | GLS | 5'UTR | 5 | 39.0 | -8130 | Reticulocyte fraction of red cells | 2 | 190872738 | rs4853525 | 0.3682 | 1.00E-09 | 9 | 0.02257269 | 27863252 | Astle WJ |
| chr2 | 190880868 | 190880920 | GCA | GLS | 5'UTR | 5 | 39.0 | -8130 | High light scatter reticulocyte percentage of red cells | 2 | 190872738 | rs4853525 | 0.3682 | 9.00E-12 | 11.04575749 | 0.02538071 | 27863252 | Astle WJ |
| chr2 | 190880868 | 190880920 | GCA | GLS | 5'UTR | 5 | 39.0 | -8130 | High light scatter reticulocyte count | 2 | 190872738 | rs4853525 | 0.3682 | 3.00E-12 | 11.52287875 | 0.02604801 | 27863252 | Astle WJ |
| chr2 | 190880868 | 190880920 | GCA | GLS | 5'UTR | 5 | 39.0 | -8130 | Reticulocyte count | 2 | 190872738 | rs4853525 | 0.3682 | 7.00E-10 | 9.15490196 | 0.02302138 | 27863252 | Astle WJ |
| chr2 | 190880868 | 190880920 | GCA | GLS | 5'UTR | 5 | 39.0 | -4397 | Immature fraction of reticulocytes | 2 | 190876471 | rs11687659 | 0.3445 | 5.00E-11 | 10.30103 | 0.02456787 | 27863252 | Astle WJ |
| chr11 | 119206289 | 119206323 | CGG | CBL | 5'UTR | 5 | 44.5 | -1676 | Plateletcrit | 11 | 119204613 | rs4938637 | 0.0742 | 5.00E-52 | 51.30103 | 0.1059334 | 27863252 | Astle WJ |
| chr11 | 119206289 | 119206323 | CGG | CBL | 5'UTR | 5 | 44.5 | 4 | Platelet count | 11 | 119206293 | rs7108857 | NR | 4.00E-34 | 33.39794001 | 0.0532 | 29403010 | Kanai M |
| chr11 | 119206289 | 119206323 | CGG | CBL | 5'UTR | 5 | 44.5 | 6319 | Platelet count | 11 | 119212608 | rs36109901 | 0.2722 | 3.00E-43 | 42.52287875 | 0.05652886 | 27863252 | Astle WJ |

**able S3**. List of nearby GWAS signal of exonic triplet repeats with high (IQR>=5) variability. GWAS studies were adopted from GWAS atalog (https://www.ebi.ac.uk/gwas/). GWAS signals within 10 kb from the start of the repeats were listed.
